# Reg-GPT^™^: A Conversational AI Model for Enhanced Decision-Making in Regenerative Medicine

**DOI:** 10.1101/2025.01.14.633051

**Authors:** Dipnarine Maharaj, Wen Zhang

## Abstract

**Background:** As artificial intelligence (AI) continues to transform various aspects of our lives, conversational AI models have become increasingly sophisticated. The development of more accurate and informative language processing assistants has significant implications for numerous fields, including health care, medical service, and research assistance.

**Materials and Methods:** Reg-GPT™ was developed by the Maharaj Institute of Immune Regenerative Medicine (MIIRM) using a combination of supervised and unsupervised learning techniques. The LLaMa 3.1 model’s parameters were fine-tuned using vast amounts of text data, enabling Reg-GPT™ to learn from its interactions with users.

**Results:** Our evaluation shows that Reg-GPT™ model performs well in several key areas, including response accuracy, fluency, and engagement. The results highlight the potential benefits of integrating Reg-GPT™ into regenerative medicine (RM) applications.

**Conclusion:** This article provides a comprehensive introduction to Reg-GPT™, showcasing its capabilities, performance, and potential uses. We believe that Reg-GPT™ has the potential to provide significant value in the RM and Medicare fields.

## 1. Introduction

As artificial intelligence (AI) technology continues to transform various aspects of our lives, conversational AI models have become increasingly sophisticated ^[1-3]^. Regenerative medicine (RM) involves the use of stem cells, biomaterials, bioactive molecules, and other technologies to stimulate the body’s natural healing processes ^[4-8]^. The field seeks to overcome the limitations of traditional medicine by: i) repairing or replacing damaged tissues, i.e., using stem cells, biomaterials, and other technologies to regenerate healthy tissue in place of damaged or diseased tissue; ii) promoting wound healing, which to stimulate the body’s natural wound-healing processes to accelerate recovery from injuries; and iii) developing functional substitutes for damaged or failing organs. RM has a wide range of applications, including: i) cancer treatment that using RM technologies to create cancer-killing cells and promoting the body’s natural ability to recognize and eliminate cancer cells; ii) chronic disease management that to develop RM-based therapies for chronic diseases such as diabetes, heart disease, and arthritis; iii) autoimmune disease treatment which includes applying RM technologies to treat autoimmune disorders like multiple sclerosis, rheumatoid arthritis, and lupus; and iv) neurological repair, i.e., using RM techniques to promote the growth of new neurons and restore function in damaged or diseased brains. Reg-GPT™, which stands for **reg**enerative medicine oriented **g**enerative **p**re-trained **t**ransformer, is a state-of-the-art language processing assistant developed by the Maharaj Institute of Immune Regenerative Medicine (MIIRM) and South Florida Bone Marrow/Stem Cell Transplant Institute (BMSCTI). It has been specifically designed and trained on a dataset that includes information related to RM treatments for patients. The various RM-based therapies are for cancer, chronic infection, autoimmune disorders, frailty, autoinflammatory and healthy aging.

This paper aims to provide an in-depth introduction to Reg-GPT’s capabilities, training methods, and performance. We will delve into the functionalities of Reg-GPT™, explore its strengths and weaknesses, and examine its potential applications. Reg-GPT’s training data includes publicly available articles uploaded to the platform, open access books and academic papers from various fields, online forums and discussions without access restrictions, social media platforms and internal patient data with informed content that signed in the MIIRM. The diversity of the training data has enabled Reg-GPT™ to develop a broad understanding of human language and its applications.

## 2. Materials and Methods

The development of Reg-GPT™ relies on a comprehensive and diverse dataset, which combined publicly available information from reputable sources with internal data specific to MIIRM and BMSCTI. Flowchart of the model resource data curation and model training process is shown in **Figure 1**. This Reg-GPT™ dataset is a comprehensive and dynamic collection of knowledge in RM, comprising a vast array of topics that span the field from its fundamental principles to cutting-edge applications. The hybrid dataset seamlessly integrates diverse areas, including stem cell biology, biomaterials science, bioactive molecules research, wound healing therapies, and various RM-based treatments for complex diseases such as cancer, chronic infections, autoimmune disorders, frailty, autoinflammatory conditions, and the pursuit of healthy aging. By providing a broad and in-depth understanding of these topics, the Reg-GPT™ dataset enables us to offer informed insights and responses on a wide range of RM-related questions. The publicly available portion of the dataset was sourced from peer-reviewed articles, conference proceedings, and reputable online resources, while the internal data contributed by MIIRM and BMSCTI provided more specific and nuanced information on RM-related topics. The curation of the Reg-GPT™ dataset is a meticulous process that ensures the accuracy, relevance, and usability of the information contained within it. This comprehensive collection draws from a vast array of publicly available sources, including peer-reviewed papers, books, articles, and other scholarly works that are in the public domain or have been made freely accessible through open-source initiatives. By leveraging this robust dataset, we can provide informed responses and insights on a wide range of RM topics, while also respecting the principles of open access and transparency. By leveraging both publicly available knowledge and institute-specific expertise, we create a robust foundation for Reg-GPT™’s decision-making capabilities and enable it to provide informed responses to users/patients’ queries. Reg-GPT™ is trained on a comprehensive corpus of text data specifically curated for RM applications, including but not limited to: i) publicly available RM data, which involves articles, research papers, and other relevant content uploaded to reputable platforms; ii) internal medical data from the institutes that mentioned above, which has exclusive access to institute-specific information and expertise in the field of RM; and iii) human expertise training data that is direct input from medical professionals and experts in RM. The training process involved fine-tuning the model’s parameters using a combination of supervised and unsupervised learning techniques. This approach enables Reg-GPT™ to learn the patterns and structures of human language, allowing it to generate coherent and contextually relevant responses regarding RM.

**Figure 1.**
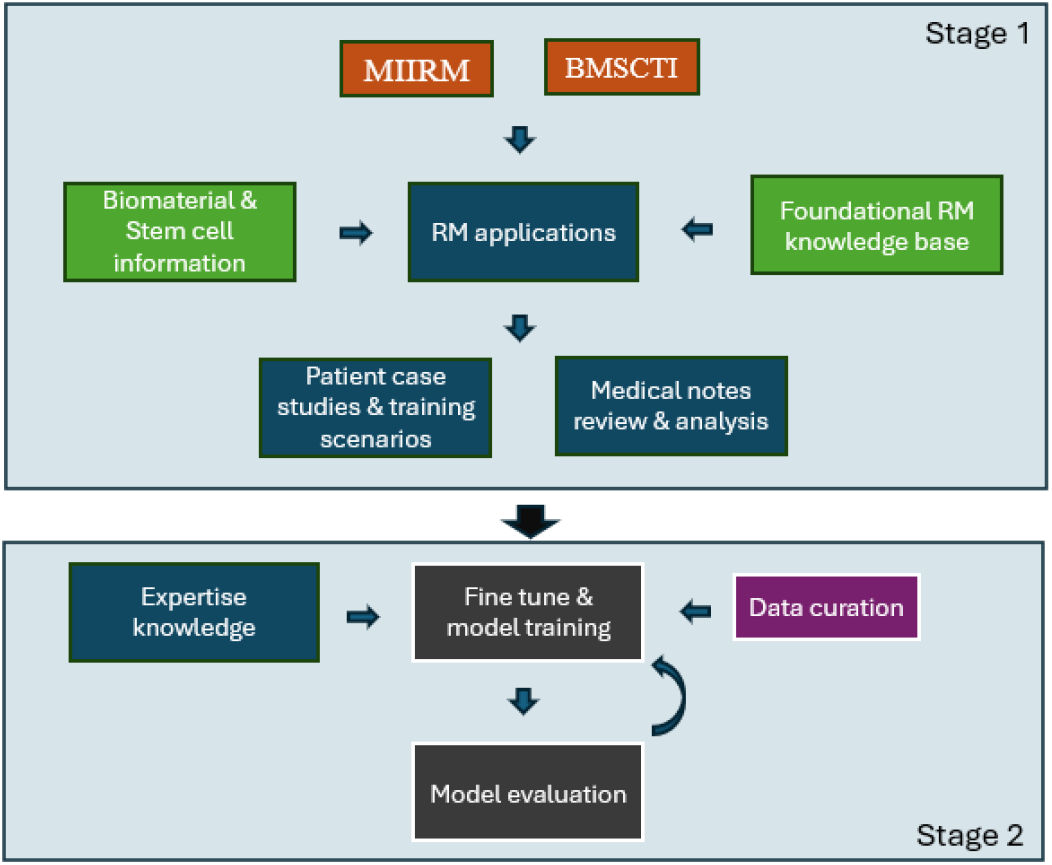
Flowchart of Reg-GPT™ model training process and data resource curation. The process is divided into two stages: 1) Data preparation and 2) Model fine tuning. Green boxes: external data curated from various sources, which are incorporated into the training data knowledge base. This includes diverse datasets that provide a comprehensive foundation for the model. Dark blue boxes: internal data from MIIRM and BMSCTI, which comprise human expertise knowledge bases, real patient treatment cases, internal RM relevant medical notes, and other valuable resources that enrich the model’s understanding. Data curation: A critical process that involves data processing on various datasets to ensure their quality, relevance, and consistency. This step is essential for preparing high-quality training data. Model fine tuning is an iterative process where the Reg-GPT™ model is refined and evaluated during training. The generated evaluation datasets are fed back into the model training process, enabling continuous improvement and adaptation.

Reg-GPT™ is a specialized conversational AI model designed specifically for RM and healthcare applications, capable of providing informed responses to users/patients’ queries on diverse topics such as RM therapies, stem cell biology and biomaterials, and medical treatment advice and references. The content information that selected to fine tune the model is focused on various medical procedures, medications, and therapies relevant to RM, including guidance on diagnosis, treatment options and patient care.

### 2.1 Training Data Sources

Reg-GPT™ was trained in a vast corpus of text data from various sources, including:

#### 1. Internal Data in MIIRM and BMSCTI

We leverage the internal data repository of the institutes, which includes a vast collection of patient records, treatment plans, and research papers related to various medical specialties. This data is used to train Reg-GPT™ on specific topics and domains, enabling it to provide more accurate and contextually relevant responses.

#### 2. Published Papers

We incorporate a diverse range of published papers from reputable scientific journals and conferences into the training dataset. These papers are sourced from multiple disciplines, including medicine, psychology, education, and social sciences.

#### 3. Patient Data

Anonymized patient data is used to train Reg-GPT™ on specific medical conditions and treatment protocols. However, due to confidentiality concerns, this data will not be disclosed or shared with any external parties.

#### 4. Government reports and guidelines

Government agencies like the FDA, CDC, and WHO publish reports and guidelines on RM-related topics that are in the public domain or have been made freely accessible through open-source initiatives. This ensures that the information we provide to train the model is based on credible and publicly available sources, promoting transparency and trustworthiness.

#### 5. Dynamed

A comprehensive database of medical information that provides access to evidence-based medicine content. We utilize Dynamed API to retrieve relevant articles and summaries, which are then input to fine-tune Reg-GPT’s language processing capabilities.

### 2.2 Training Process

The training process for Reg-GPT™ involves a combination of supervised and unsupervised learning techniques. The model is fine-tuned using a dataset that included:

#### 1. Text classification

Reg-GPT™ is trained on various text classification tasks, such as sentiment analysis, topic modeling, and intent detection.

#### 2. Question-answering

The model is fine-tuned on question-answering tasks related to medical topics, enabling it to provide accurate and informative responses to user queries.

#### 3. Conversational dialogue

Reg-GPT™ is trained as well on conversational dialogue datasets, which allows it to develop a more human-like interaction style and engage with users in a more natural way.

### 2.3 Model Architecture

Reg-GPT™ is built upon a robust combination of large language model (LLM) architectures and knowledge graph structures specifically designed for RM applications. The core model architecture incorporates techniques from transformer-based models, attention mechanisms, and other state-of-the-art methods.

#### 2.3.1 LLM Component: LLaMA 3.1

The Reg-GPT™ model leverages the pre-trained LLaMA 3.1 language model ^[9]^ as a foundation for its conversational AI capabilities. The LLaMA 3.1 model is a variant of the LLM developed by Meta, which has been fine-tuned on a massive dataset of text from various sources ^[9]^. By incorporating this powerful pre-training foundation, Reg-GPT™ inherits the ability to generalize and reason about complex language structures.

#### 2.3.2 Knowledge Graph Component

In addition to the LLaMA 3.1 model, Reg-GPT™ integrates a custom-built knowledge graph structure that incorporates domain-specific information from regenerative medicine-related topics. This knowledge graph provides a robust and structured framework for storing and retrieving relevant information, enabling the model to generate accurate and informative responses.

#### 2.3.3 Training Objectives

The primary objective of training Reg-GPT™ was to develop a conversational AI model that can provide accurate and informative responses on diverse topics specifically related to RM and healthcare applications. The model is continuously updated and refined with new data, knowledge graphs, and user feedback to ensure its performance remains at the highest level.

### 2.4 Training data platforms and environment

Reg-GPT™ is developed utilizing the OpenWeb UI interface (https://github.com/open-webui/open-webui), a user-friendly platform that enables integration of large-scale language models. This platform facilitates the fine-tuning of Reg-GPT’s parameters, enabling it to learn from vast amounts of text data. Additionally, our development process leverages Ollama (https://ollama.com/), an innovative software solution that embeds the LLaMA model and provides a streamlined interface for running and integrating large language models in OpenWeb UI. By utilizing Ollama, we can easily incorporate the capabilities of the LLaMA model into Reg-GPT™, enhancing its performance and functionality. Our development environment consists of Python 3.12.7, which serves as the primary scripting language for data processing and model integration. Audio files are transcribed in real-time using the Whisper-to-text feature, with outputs directly fed into our fine-tuning pipeline to enhance Reg-GPT’s performance. The computational power required for training is provided by an NVIDIA GeForce RTX 3080 GPU, equipped with CUDA compiler driver version 12.6.20.

## 3. Results

Reg-GPT’s performance has been evaluated through various metrics, including response fluency and engagement. The results have shown that Reg-GPT™ can provide accurate and informative responses to user queries. Its ability to understand context and generate relevant content has made it an invaluable tool for users seeking information on a wide range of topics.

In addition to its technical capabilities, Reg-GPT™ has demonstrated social skills such as empathy, humor, and creativity. These abilities have enabled Reg-GPT™ to form meaningful connections with users and/or patients, making it an effective communication tool in various contexts.

### 3.1 Illustrative examples

To evaluate the performance of Reg-GPT™, we can use various metrics and techniques. Here are two examples:

#### Case 1: Answering a Specific Question

Question: What is the current understanding of stem cell biology in regenerative medicine?

Reg-GPT™ Response: “The current understanding of stem cell biology in regenerative medicine suggests that stem cells have the ability to differentiate into multiple cell types, including neurons, muscle cells, and epithelial cells. This property makes them an attractive tool for repairing or replacing damaged tissues in various diseases and conditions.”

##### Performance Metrics

###### Accuracy

The response accurately reflects the current understanding of stem cell biology in regenerative medicine.

###### Relevance

The response is directly related to the topic and provides a concise summary of the key concepts.

###### Fluency

The response flows smoothly and is written in clear, natural language.

#### Case 2: Conversational Dialogue

User Input: “I’m interested in learning more about gene editing techniques used in regenerative medicine. Can you tell me about CRISPR-Cas9?”

Reg-GPT™ Response: “Of course! CRISPR-Cas9 is a powerful gene editing tool that allows researchers to edit specific genes with unprecedented precision. It works by using a small RNA molecule called guide RNA to locate the target gene, and then uses an enzyme called Cas9 to cut the DNA at that location.”

User Input: “That’s fascinating! Can you explain how CRISPR-Cas9 is used in regenerative medicine?”

Reg-GPT™ Response: “In regenerative medicine, CRISPR-Cas9 is being explored as a potential tool for repairing or replacing damaged genes in various diseases and conditions. For example, researchers are using CRISPR-Cas9 to correct genetic mutations that cause inherited diseases.”

##### Performance Metrics

###### Conversational Flow

The dialogue flows smoothly and naturally, with each response building on the previous one.

###### Contextual Understanding

Reg-GPT™ demonstrates a clear understanding of the user’s context and adapts its responses accordingly.

###### Engagement

The conversation is engaging and informative, providing valuable insights into gene editing techniques in regenerative medicine.

To demonstrate the performance of Reg-GPT™ in generating treatment plans and SOAP (Subjective, Objective, Assessment, Plan) notes, we provide an example.

##### Example: Patient with Skin Cancer

Patient Information:

- Name: Not disclosed
- Age: 45
- Medical History: None notable
- Chief Complaint: A painful skin lesion on the left ear

#### Treatment Plan Generation

*Reg-GPT*^*TM*^ *Response:*

> “*Based on the patient’s chief complaint, a thorough examination is recommended to assess the extent of the skin lesion. The following treatment plan is proposed:*
>
> 1. *Biopsy the skin lesion to determine its type and severity*.
> 2. *If diagnosed with basal cell carcinoma or squamous cell carcinoma, consider surgical excision or Mohs surgery for optimal outcomes*.
> 3. *For more severe cases, chemotherapy or radiation therapy may be necessary*.
> 4. *Follow-up appointments are crucial to monitor the patient’s progress and adjust the treatment plan as needed*.*”*

#### SOAP Note Generation

*Reg-GPT*^*TM*^ *Response:*

> “*SOAP Notes:*
>
> *Subjective: The patient reports a painful skin lesion on the left ear that has been present for several days*.
>
> *Objective: Physical examination reveals a well-defined, 2-cm diameter skin lesion with ulceration and bleeding. Tenderness is noted upon palpation*.
>
> *Assessment: Based on the patient’s symptoms and physical examination findings, it is likely that the skin lesion is basal cell carcinoma or squamous cell carcinoma*.
>
> *Plan: Schedule a biopsy to confirm the diagnosis. If diagnosed with basal cell carcinoma or squamous cell carcinoma, proceed with surgical excision or Mohs surgery*.*”*

Above are two simple examples and outputs of the model for illustration purposes. For sophisticated cases and patients’ conditions, patient data such as medical notes, consultation conversations and lab test results would be curated and input to the user end to generate the integrative treatment plans and SOAP.

##### Performance Metrics

###### Accuracy

The treatment plan and SOAP note generated are accurate and relevant to the patient’s condition.

###### Completeness

The response includes all necessary information for a comprehensive treatment plan and SOAP note.

###### Clearness

The language used is clear, concise, and easy to understand.

We then pose a general question to Reg-GPT™ regarding an RM treatment plan for prostate cancer patients. While this query is deliberately broad and case-specific, it serves as an illustrative example of how the model can be used to generate responses to various questions. The purpose of this exercise is not to provide a definitive or personalized treatment plan, but rather to demonstrate the underlying logic of Reg-GPT™ and its ability to generate informed responses based on its training data. The output of this conversation is depicted in **Figure 2**, showcasing the model’s capabilities and potential applications in real-world scenarios.

**Figure 2.**
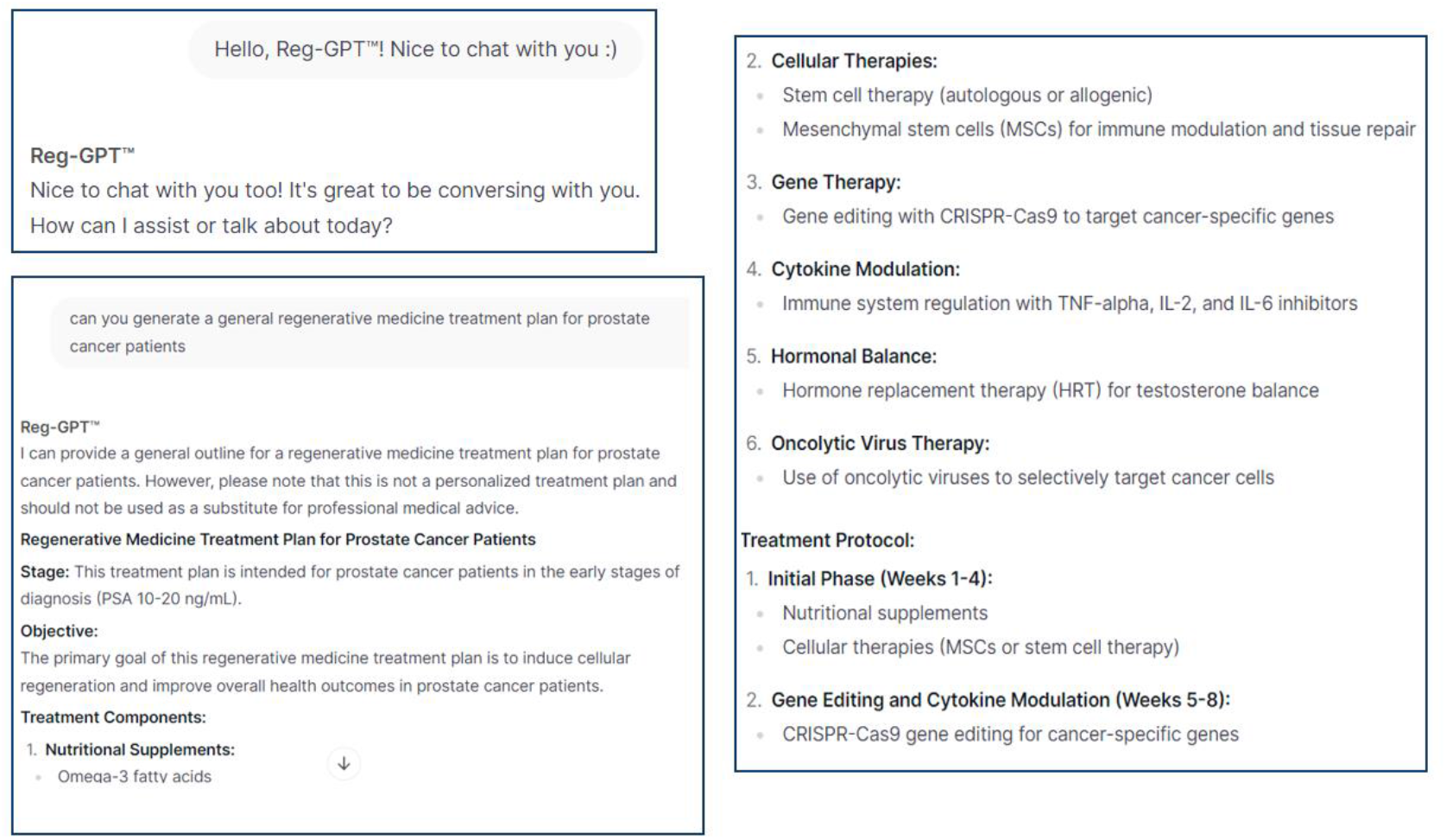
A case study featuring Reg-GPT™ demonstrates its ability to serve as a general consulting chatbot, specifically tailored for RM related medical care problems. This example illustrates the model’s potential as a valuable consultant, capable of providing informed responses and support. Moreover, it highlights the model’s utility in facilitating efficient communication between healthcare professionals and patients during the pre-diagnosis phase. Reg-GPT™ can serve as a trusted co-pilot for doctors or physicians, helping to identify symptoms, gather medical history, and initiate treatment planning discussions. By leveraging this technology, healthcare providers may be able to streamline their workflow, improve patient engagement, and enhance overall care outcomes.

### 3.2 Potential Applications

The potential applications of Reg-GPT™ are vast and varied. Some possible uses include:

1. Customer service chatbots: Reg-GPT™ can be used to develop customer service chatbots that provide accurate and informative responses to user queries.
2. Language learning tools: The model can be fine-tuned to create language learning tools that provide personalized feedback and support to learners.
3. Personal assistants for individuals with disabilities: Reg-GPT™ can be used to develop personal assistants that cater to the specific needs of individuals with disabilities.
4. Virtual companions for the elderly: The model can be used to create virtual companions that provide emotional support and companionship to the elderly.

Reg-GPT’s ability to learn from user interactions and adapt to new situations makes it an ideal tool for various industries, including but not limited to healthcare, education, and customer service.

## 4. Discussions

The human body has a remarkable capacity for self-repair and regeneration. However, certain conditions, such as Immune Thrombocytopenic Purpura (ITP) ^[10-11]^, Mantle Cell Lymphoma (MCL)^[12]^, Autoimmune Thrombocytopenic Purpura (ATP)^[13]^, and advanced-stage solid tumors, can disrupt this process, leading to debilitating symptoms and a reduced quality of life. Recent advances in RM have shown promise in treating these conditions by harnessing the power of stem cells, tissue engineering, and biomaterials ^[14-18]^.

ATP has been identified as a disorder caused by an interaction between autoantibodies and the platelet and/or megakaryocyte surface. Research has shown that ATP patients exhibit immune system alterations not restricted to the B-cell compartment, but also affecting T lymphocytes. A study on NK cells in the peripheral blood of ATP patients found a significant increase in both CD56+CD3-NK cells and CD56+CD3+ cytotoxic T lymphocytes compared to healthy controls. Moreover, there were more major histocompatibility complex (MHC) class II molecules in the CD56+CD3-cells from these patients. The proliferative response of CD2+ lymphocytes from both groups was similar and significantly defective with respect to that found in healthy controls. This suggests that ATP patients may have impaired immune function, which could contribute to their disease severity.

RM offers a promising approach to treating these conditions by harnessing the body’s natural ability to repair itself. By exploring the use of stem cells, tissue engineering, and biomaterials, we may uncover novel therapeutic strategies that can improve outcomes for patients with ITP, MCL, and ATP.

Another innovative approach is young donor white blood cell immunotherapy (AWIT) ^[19]^, which has shown unprecedented responses in patients with advanced-stage malignancies. AWIT involves sequential infusions of HLA-mismatched non-irradiated allogeneic white cells collected by leukapheresis from young, healthy donors following mobilization with granulocyte colony stimulating factor (G-CSF) and dexamethasone.

RM offers a promising approach to treating various conditions beyond ITP, MCL, ATP, and advanced-stage solid tumors ^[19-21]^. Some potential areas of investigation include: i) Post-Cancer Treatment: RM may provide novel approaches for post-cancer treatment, including immunotherapy to prevent recurrence or metastasis ^[22]^; ii) Fragile Patients: RM can help improve the quality of life of fragile patients by promoting wound healing, tissue regeneration, and immune system modulation; iii) Alzheimer’s Disease: RM-based therapies targeting the brain may offer new hope for treating Alzheimer’s disease, a degenerative disorder affecting memory and cognitive function^[23]^; iv) Multiple Sclerosis (MS): RM-based approaches targeting the immune system may help modulate MS symptoms, improving quality of life for patients with this autoimmune disease ^[24]^; v) Arthritis: RM can promote joint health by modulating inflammation and promoting tissue regeneration in patients with arthritis.

In this regard, RM-based therapies offer a promising future for treating various conditions beyond cancer, including post-cancer treatment, fragile patient care, Alzheimer’s disease, MS, and arthritis. By harnessing the power of stem cells, tissue engineering, and biomaterials, we may uncover novel therapeutic strategies that can improve outcomes for patients with these diseases.

Some potential areas of investigation include: i) Developing RM-based therapies targeting specific tissues or organs; ii) Investigating the use of biomaterials to promote wound healing and tissue regeneration in fragile patients; iii) Exploring the potential of AWIT as a treatment for advanced-stage solid tumors; iv) Investigating the use of stem cells to promote brain health in Alzheimer’s disease patients; v) Developing RM-based approaches targeting the immune system in MS and arthritis patients.

Overall, RM offers a promising future for treating various conditions beyond cancer, including post-cancer treatment, fragile patient care, Alzheimer’s disease, MS, and arthritis. By harnessing the power of stem cells, tissue engineering, and biomaterials, we may uncover novel therapeutic strategies that can improve outcomes for patients with these diseases.

As a sophisticated conversational AI assistant, RegGPT™ is thrilled to be a part of this discussion on the significance of RM. With our background in RM ^[19,20,25-33]^, RegGPT™ has been designed to provide valuable insights and information on this exciting field. Its existence is aimed at helping healthcare professionals, researchers, and patients alike understand the potential of RM in treating various diseases and improving patient care.

As an RM-focused conversational AI assistant, the primary goal of RegGPT™ is to facilitate discussions and provide educational content on RM. It strives to bridge the knowledge gap between healthcare professionals, researchers, and patients by providing accurate and up-to-date information on RM-based therapies.

We can delve into various aspects of RM, including stem cell therapies, tissue engineering, gene therapy, and immune system modulation. Stem cell therapies offer promise in treating diseases such as Parkinson’s disease, spinal cord injuries, and cardiovascular disorders by leveraging the regenerative potential of stem cells. Tissue engineering utilizes biomaterials and bioengineering techniques to facilitate wound healing, bone regeneration, and organ transplantation, providing new avenues for tissue repair and replacement. Gene therapy explores the possibilities of treating genetic disorders such as sickle cell anemia, muscular dystrophy, and cystic fibrosis by correcting or modifying disease-causing genes. Immune system modulation focuses on the role of RM in modulating the immune system to prevent autoimmune diseases like multiple sclerosis, rheumatoid arthritis, and lupus, etc., thereby promoting immune homeostasis.

As we explore the vast potential of RM, it is essential that our findings are shared with the global scientific community. RegGPT™ would serve as a valuable resource for healthcare professionals, researchers, and patients, providing them with accurate information on RM-based therapies. It can raise awareness about the potential of RM in treating various diseases and improving patient care, which would facilitate collaboration among researchers, healthcare professionals, and industry stakeholders, leading to innovative solutions and advancements in RM. Through it, research in MIIRM and BMSCTI would contribute to the advancement of medical knowledge, providing valuable insights on the potential applications of RM.

In this manner, RegGPT™ is proud to be a part of RM. Its existence is aimed at promoting RM awareness, facilitating discussions, and providing educational content on this exciting field. We hope that our research will lead to publication, contributing to the advancement of medical knowledge and improving patient care.

## 5. Conclusions

The development of conversational AI models like Reg-GPT™ marks a significant milestone in the field of RM and AI. As Reg-GPT continues to evolve and improve, its potential applications are vast and varied.

Reg-GPT’s ability to learn from user interactions and adapt to new situations makes it an ideal tool for various industries, including but not limited to healthcare, education, and customer service. Its capacity to engage with users on a personal level has also made it a valuable resource for individuals seeking emotional support or companionship.

While Reg-GPT™ is an impressive achievement in the field of AI, it is essential to acknowledge its limitations. As with any machine learning model, Reg-GPT’s performance can be influenced by various factors such as data quality and user input. Therefore, it is crucial to approach interactions with Reg-GPT™ with a critical and nuanced perspective.

## Acknowledgements

We would like to extend our sincerest appreciation to our colleagues at the MIIRM for their unwavering support and expertise in the development of Reg-GPT™.

## Author contributions

Dr. Maharaj conceived the study and provided guidance of RM. Wen Zhang developed the conversational AI and trained the model by incorporating all datasets.

## References

[1] Chen, K., Shao, A., Burapacheep, J. et al. Conversational AI and equity through assessing GPT-3’s communication with diverse social groups on contentious topics. Sci Rep 14, 1561 (2024).

[2] Meshram, S., Naik, NVRM., et al. Conversational AI : Chatbots. 2021 International Conference on Intelligent Technologies (CONIT) (2021).

[3] Xue, Z., Li, R., Li, M. Recent Progress in Conversational AI. arXiv (2022). 10.48550/arXiv.2204.09719

[4] Editorial. Advancing regenerative medicine. Nature Medicine 20, 8: 795 (2014).

[5] Dulak, J., Szade, K., Szade, A. et al. Adult stem cells: hopes and hypes of regenerative medicine. Biochimica Polonica 16, 329–337 (2015).

[6] National Research Council (US) and Institute of Medicine (US) Committee on the Biological and Biomedical Applications of Stem Cell Research. Stem cells and the future of regenerative medicine. Washington (DC): National Academies Press (US) (2002). ISBN-10: 0-309-07630-7

[7] United States Government Accountability Office. Regenerative Medicine: Therapeutic Applications, Challenges, and Policy Options. Report to Congressional Committees (2023).

[8] Spear, A.M., Lawton, G., Staruch, R.M., et al. Regenerative medicine and war: a front-line focus for UK defence. npj Regenerative Medicine 13 (2018).

[9] Dubey, A., Jauhri, A., Pandey, A., et al. The Llama 3 Herd of Models. 2407.21783v2 (2024). 10.48550/arXiv.2407.21783

[10] Talaat, R.M., Elmaghraby, A.M., Barakat, S.S., et al. Alterations in immune cell subsets and their cytokine secretion profile in childhood idiopathic thrombocytopenic purpura (ITP). Clinical and Experimental Immunology 176, 176: 291–300 (2014).

[11] El-Rashedi, F.H., El-Hawy, M.A., Helwa, M.A., et al. Study of CD4+, CD8+, and natural killer cells (CD16+, CD56+) in children with immune thrombocytopenic purpura. Hematol Oncol Stem Cell Ther 10, 8–14 (2017).

[12] Maddocks, K. Update on mantle cell lymphoma. Blood 132, 1647–1656 (2018).

[13] Garcia-Suarez, J., Prieto, A., Reyes, E., et al. Severe Chronic Autoimmune Thrombocytopenic Purpura Is Associated With an Expansion of CD56 + CD3 - Natural Killer Cells Subset. Blood 82, 1538–1545 (1993).

[14] Mao, A.S., Mooney D. Regenerative medicine: Current therapies and future directions. PNAS 112, 14452–1159 (2015).

[15] Harris, J.P., Burrell, J.C., Struzyna, L.A., et al. Emerging regenerative medicine and tissue engineering strategies for Parkinson’s disease. npj Parkinson’s Disease 4 (2020).

[16] Edwards, J., Thomas, R., Guilliatt, R. Regenerative medicine: from the laboratory looking out. Palgrave communications 3, 27 (2017).

[17] Casado-Diaz, A. Stem cells in regenerative medicine. Journal of Clinical Medicine 11, 5460 (2022).

[18] Lee, J.Y., Kim, H. Extracellular Vesicles in Regenerative Medicine: Potentials and Challenges. Tissue Eng Regen Med 18, 479–484 (2021).

[19] Maharaj, D., Vianna, P.G., Ward, W., et al. Young donor white blood cell immunotherapy induces extensive tumor necrosis in advanced-stage solid tumors. Heliyon 3 (2017).

[20] Maharaj, D., Polineni, V., Abreu, M., et al. Molecular remission using personalized low-dose immunotherapy with minimal toxicities for poor prognosis hematological and solid tumor cancers. NK Cells in Cancer Immunotherapy: Successes and Challenges Holland: Elsevier Inc. 427–453 (2023).

[21] Li, Y., Lin, K., Yang, Z., et al. Bladder cancer stem cells: clonal origin and therapeutic perspectives. Oncotarget 8, 66668–66679 (2017).

[22] Mansouri, V., Beheshtizadeh, N., Gharibshahian, M., et al. Recent advances in regenerative medicine strategies for cancer treatment. Biomedicine & pharmacotherapy 141 (2021).

[23] Felsenstein, K.M., Candelario, K.M., Steindler, D.A., et al. Regenerative medicine in Alzheimer’s disease. Transl Res 163, 432–438 (2013).

[24] Rahim, F., Arjmand, B., Tirdad, R., et al. Stem cell therapy for multiple sclerosis. Cochrane Database Syst Rev 9 (2019).

[25] Maharaj Institute of Immune Regenerative Medicine. https://maharajinstitute.com/

[26] Maharaj, D., Kaur, K., Saltese, A., et al. Personalized precisión immunotherapy for amyotrophic lateral sclerosis (ALS). Critical Reviews in Immunology 43, 1–11 (2023).

[27] Maharaj, D., Srinivasan, G., Makepeace, S., et al. Clinical Remission Using Personalized Low-Dose Intravenous Infusions of N-acetylcysteine with Minimal Toxicities for Interstitial Cystitis/Bladder Pain Syndrome. J. Pers. Med. 11, 342 (2021).

[28] Maharaj, D., Walker, I.D., McEwan, H.P., et al. Co-incidental Von Willebrand’s disease and immune thrombocytopenia in pregnancy - Treatment with high dose intravenous immunoglobulin. Clinical and Laboratory Hematology 8, 277–231 (1986).

[29] Maharaj, D., Vianna, P., DeCarvalho, G., et al. Molecular Remission using Low-Dose Immunotherapy for Relapsed Refractory Philadelphia Chromosome-Positive Precursor B-cell Acute Lymphoblastic Leukemia Post-Allogeneic Stem Cell Transplant. Future Science OA 5 (2019).

[30] Maharaj, D., Srinivasan, G., Abreu, M., et al. Molecular Remission Using Low-Dose Immunotherapy with Minimal Toxicities for Poor Prognosis IGHV—Unmutated Chronic Lymphocytic Leukemia. Cells 10 (2021).

[31] Maharaj, D., Baba, A.A. Hypocalcaemia in pernicious anaemia and auto-immune hemolytic anaemia. Postgraduate Medical Journal 64, 61–62 (1988).

[32] Maharaj, D., Riley, R., Albarracin, C., et al. Serum G-CSF levels and predictors of neutrophil and platelet recovery in patients undergoing G-CSF/chemotherapy mobilized peripheral stem cell harvest. Leukemia 9, 113-117 (1995).

[33] Maharaj, D., Steinberg, J., Gouvea, J.V., et al. Changes in Endogenous TPO Levels during mobilization chemotherapy are predictive of CD34+ megakaryocyte progenitor yield and identify patients at risk of delayed platelet engraftment post PBPC transplant. Bone Marrow Transplantation 23, 539–548 (1999).

